## Supplementary data for "BiSCoT: Improving large eukaryotic genome assemblies with optical maps"

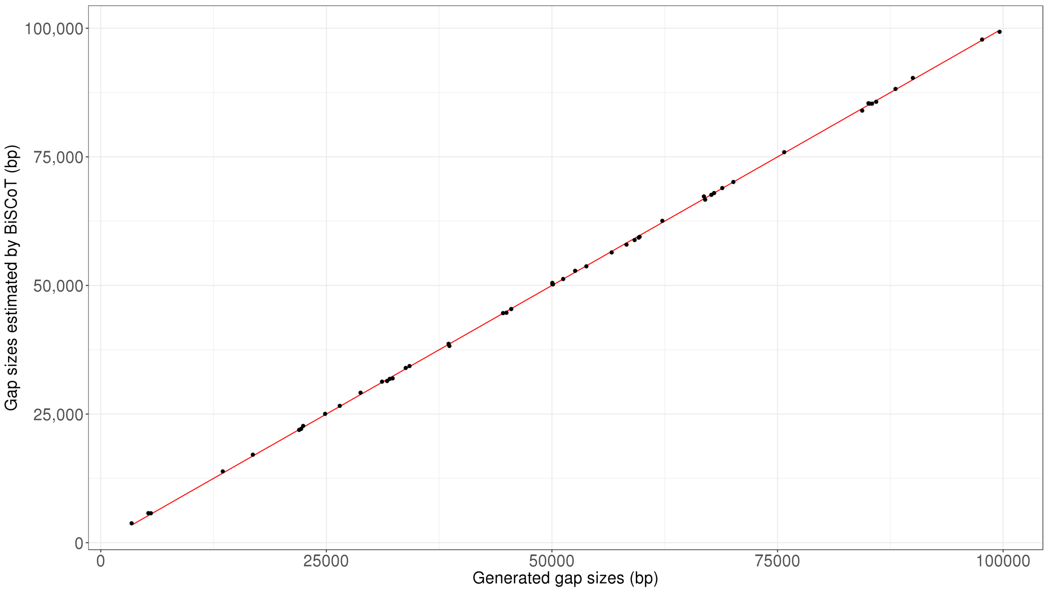


**Supplementary Figure 1.** Distribution of gap sizes estimated by BiSCoT using the optical maps against the gap sizes introduced in the simulated Human chromosome 1 assembly

| | Overlaps | Gaps | Contained contigs |
|---|---|---|---|
| Generated number of events | 50 | 50 | 5 |
| Cumulative size of events before BiSCoT | 2,206,983 | 2,547,879 | 1,168,275 |
| Mean size of events before BiSCoT | 44,139 | 50,957 | 233,655 |
| Min size of events before BiSCoT | 278 | 3,420 | 215,581 |
| Max size of events before BiSCoT | 98,683 | 99,611 | 283,879 |
| Remaining number of events after BiSCoT | 11 | 50 | 0 |
| Cumulative size of events after BiSCoT | 30,587 | 2,549,286 | - |
| Mean size of events after BiSCoT | 2680 | 50,985 | - |
| Min size of events after BiSCoT | 278 | 3,392 | - |
| Max size of events after BiSCoT | 5,959 | 99,792 | - |

**Supplementary Table 1.** Metrics of the overlaps, gaps and contained contigs introduced in the simulated Human genome's chromosome 1 assembly before and after applying BiSCoT.

| | Simulated contigs | Bionano | | BiSCoT | |
|---|---|---|---|---|---|
| | | Contigs | Scaffolds | Contigs | Scaffolds |
| Cumulative size | 231,215,374 | 231,215,374 | 252,023,317 | 227,870,703 | 248,278,703 |
| N50 | 2,429,486 | 2,363,641 | 62,178,504 | **3,612,465** | 61,255,664 |
| L50 | 46 | 43 | 2 | **19** | 2 |
| N90 | **1,677,340** | 1,621,176 | 2,792,747 | 1,658,986 | 2,872,288 |
| L90 | 95 | 90 | 10 | **55** | 9 |
| auN | 2,347,866 | 2,260,168 | 62,408,635 | **4,569,037** | 62,312,336 |
| # Ns | 0 | 0 | 20,789,288 | 0 | 20,408,000 |
| NGA50 | 2,354,935 | 2,287,164 | 36,067,187 | **2,931,345** | 42,227,440 |
| NGA75 | 1,943,500 | 1,803,364 | 2,537,018 | **2,034,390** | 7,636,847 |
| # misassemblies | **0** | **0** | 38 | **0** | 12 |

**Supplementary Table 2.** Metrics of the simulated contigs of the NA12878 chromosome scaffolds and contigs before or after BiSCoT treatment. Bold formatting indicates the best scoring assembly among contigs.

| | Nanopore contigs | Bionano | | BiSCoT | |
|---|---|---|---|---|---|
| | | Contigs | Scaffolds | Contigs | Scaffolds |
| Cumulative size | 547,048,580 | 547,048,579 | 557,059,506 | 544,091,840 | 553,538,342 |
| N50 | 7,294,619 | 6,977,344 | 29,588,784 | **12,590,381** | 29,458,880 |
| L50 | 23 | 23 | 8 | **15** | 8 |
| N90 | 1,304,967 | 1,183,704 | 13,953,050 | **2,419,034** | 13,949,143 |
| L90 | 85 | 88 | 17 | **52** | 17 |
| auN | 9,034,610 | 8,946,525 | 29,924,521 | **12,646,724** | 29,817,651 |
| # Ns | 0 | 0 | 10,010,927 | 0 | 9,446,502 |
| Complete BUSCOs | **1,601 (99.2%)** | **1,601 (99.2%)** | 1,598 (99.0%) | **1,601 (99.2%)** | 1,600 (99.2%) |
| Duplicated BUSCOs | 235 (14.6%) | 235 (14.6%) | 232 (14.4%) | **234 (14.5%)** | 232 (14.4%) |
| Missing BUSCOs | 10 (0.6%) | **9 (0.6%)** | 10 (0.6%) | **9 (0.6%)** | 10 (0.6%) |

**Supplementary Table 3.** Metrics of the *Brassica oleracea* HDEM scaffolds and contigs before or after BiSCoT treatment. Bold formatting indicates the best scoring assembly among contigs.

| | Nanopore contigs | Bionano | | BiSCoT | |
|---|---|---|---|---|---|
| | | Contigs | Scaffolds | Contigs | Scaffolds |
| Cumulative size | 373,437,357 | 373,437,357 | 406,471,180 | 369,747,840 | 402,627,824 |
| N50 | 3,793,063 | 3,603,274 | 15,479,745 | **5,519,975** | 15,275,286 |
| L50 | 25 | 26 | 8 | **17** | 8 |
| N90 | **202,023** | 154,330 | 1,748,645 | 181,213 | 1,674,920 |
| L90 | 264 | 309 | 31 | **221** | 31 |
| auN | 5,532,997 | 5,453,354 | 18,883,302 | **7,237,727** | 18,700,050 |
| # Ns | 0 | 0 | 33,033,823 | 0 | 32,879,984 |
| Complete BUSCOs | 1,604 (99.4%) | **1,605 (99.5%)** | 1,604 (99.4%) | **1,605 (99.5%)** | 1,604 (99.4%) |
| Duplicated BUSCOs | **233 (14.4%)** | 235 (14.6%) | 234 (14.5%) | **233 (14.4%)** | 233 (14.4%) |
| Missing BUSCOs | **7 (0.5%)** | **7 (0.5%)** | 7 (0.5%) | **7 (0.5%)** | 7 (0.5%) |

**Supplementary Table 4.** Metrics of the *Brassica rapa* Z1 scaffolds and contigs before or after BiSCoT treatment. Bold formatting indicates the best scoring assembly among contigs.

| | Nanopore contigs | Bionano | | BiSCoT | |
|---|---|---|---|---|---|
| | | Contigs | Scaffolds | Contigs | Scaffolds |
| Cumulative size | 518,619,765 | 518,619,765 | 526,521,784 | 517,940,161 | 525,719,686 |
| N50 | 4,019,832 | 2,097,979 | 36,762,080 | **7,987,169** | 36,858,856 |
| L50 | 33 | 60 | 6 | **24** | 6 |
| N90 | 554,125 | 292,444 | 9,697,206 | **888,370** | 9,721,221 |
| L90 | 180 | 310 | 15 | **92** | 15 |
| auN | 5,390,023 | 5,285,943 | 33,951,065 | **7,477,787** | 33,460,868 |
| # Ns | 0 | 0 | 7,902,019 | 0 | 7,779,525 |
| Complete BUSCOs | 1,558 (96.6%) | **1,562 (96.8%)** | 1,561 (96.7%) | 1,560 (96.6%) | 1,559 (96.6%) |
| Duplicated BUSCOs | 69 (4.3%) | **68 (4.2%)** | 68 (4.2%) | **68 (4.2%)** | 70 (4.3%) |
| Missing BUSCOs | **34 (2.0%)** | **34 (2.0%)** | 34 (2.0%) | **34 (2.0%)** | 34 (2.0%) |

**Supplementary Table 5.** Metrics of the *Musa schizocarpa* scaffolds and contigs before or after BiSCoT treatment. Bold formatting indicates the best scoring assembly among contigs.

| | Nanopore contigs | Bionano | | BiSCoT | |
|---|---|---|---|---|---|
| | | Contigs | Scaffolds | Contigs | Scaffolds |
| Cumulative size | 652,555,937 | 652,555,937 | 665,966,510 | 649,440,360 | 662,857,763 |
| N50 | 2,985,938 | 2,985,938 | 31,920,664 | **3,969,296** | 31,819,818 |
| L50 | 51 | 51 | 10 | **42** | 10 |
| N90 | 488,936 | 485,536 | 13,186,102 | **612,779** | 13,076,771 |
| L90 | 267 | 261 | 21 | **216** | 21 |
| auN | 4,975,870 | 4,969,973 | 29,298,774 | **5,711,235** | 29,207,200 |
| # Ns | 0 | 0 | 13,410,573 | 0 | 13,417,403 |
| Complete BUSCOs | 1,569 (97.3%) | 1,572 (97.4%) | 1,576 (97.6%) | **1,576 (97.6%)** | 1,573 (97.4%) |
| Duplicated BUSCOs | **30 (1.9%)** | 31 (1.9%) | 31 (1.9%) | 31 (1.9%) | 31 (1.9%) |
| Missing BUSCOs | 26 (1.5%) | 24 (1.5%) | 23 (1.5%) | **23 (1.5%)** | 24 (1.5%) |

**Supplementary Table 6.** Metrics of the *Sorghum bicolor Tx430* scaffolds and contigs before or after BiSCoT treatment. Bold formatting indicates the best scoring assembly among contigs.
